# Pluripotency Factors Modulate Interferon Signaling in Embryonic Stem Cells

**DOI:** 10.64898/2026.03.23.713714

**Authors:** Qing Yang, Monica Padilla-Galvez, Skyler Uhl, Julie Eggenberger, Sophie Kogut, Sara Becker, Shuibing Chen, Brad R. Rosenberg, Daniel Blanco-Melo

## Abstract

Despite lacking a robust interferon response, pluripotent stem cells remain highly resistant to viral infection, in part through the constitutive expression of immune genes traditionally classified as interferon-stimulated genes. While interferon signaling has been shown to be incompatible with the maintenance of pluripotency, the molecular mechanisms underlying this relationship remain poorly understood. Here, we investigate the transcriptional response of human embryonic stem cells (hESCs) to infection with a potent activator of the interferon response, an influenza A virus mutant lacking the viral NS1 protein. Single-cell RNA sequencing revealed that while most hESCs remain unresponsive to infection, a distinct subpopulation expresses type I and III interferons. Notably, only interferon-expressing cells mounted a robust antiviral response, characterized by strong induction of interferon-stimulated genes. In contrast to the bulk hESC population, interferon responding cells exhibited reduced expression of core pluripotency factors as well as negative regulators of interferon signaling, such as SOCS1 and SPRY4. Depletion of SOCS1 enabled hESCs to respond robustly to interferon stimulation, showing that this negative regulator is a key suppressor of interferon signaling in pluripotent stem cells. We further show that SOCS1 and additional negative regulators of IFN signaling are intrinsically expressed in hESCs and are transcriptionally controlled by pluripotency factors, such as NANOG, SOX2 and OCT4. Together, our findings support a model in which pluripotency factors regulate intrinsic immune gene expression, including negative regulators of interferon signaling, thereby suppressing canonical interferon signaling to preserve pluripotency while maintaining antiviral resistance.

**IMPORTANCE:** By combining single-cell transcriptomics with functional studies, we demonstrate that the pluripotency transcriptional program active in pluripotent stem cells coordinately regulates pluripotency factors, antiviral genes, and negative regulators of interferon signaling. This integrated control enables pluripotent stem cells to achieve effective protection against viral infection while preserving their differentiation potential, providing new insights into how innate immunity is selectively constrained in pluripotent stem cells. These findings have important implications for stem cell–based therapies, where transient modulation of antiviral responses without disrupting pluripotency could improve therapeutic efficacy. More broadly, this work advances our understanding of interferon regulation that could inform the development of antiviral strategies that enhance protective immune responses while limiting harmful or unwanted inflammatory signaling.

## INTRODUCTION

The immune system in vertebrates is built upon a complex network of responses aimed at detecting and neutralizing viral threats. Central to this defense is the interferon (IFN) response, which serves as an immediate barrier to infection and a potent activator of adaptive immunity. The IFN response is triggered when viral components, such as double-stranded RNA, are recognized by pattern recognition receptors (PRRs) such as RIG-I and MDA5 (1). This recognition initiates a cascade of intracellular signaling through key adaptor molecules, such as MAVS, which culminates in the activation of transcription factors including IRF3, IRF7, and NF-κB. These transcription factors drive the expression of Type I and Type III IFNs, which are secreted and act in autocrine and paracrine fashion (2). IFNs bind to their cognate receptors on producer and neighboring cells, activating the JAK-STAT pathway, which induces the expression of hundreds of interferon-stimulated genes (ISGs) (3). These ISGs create an antiviral state by limiting viral replication, enhancing antigen presentation, and recruiting immune cells to the site of infection (4).

In somatic cells, the IFN-mediated response is well characterized, with a clear sequence of pathogen detection, signaling, and host gene activation leading to robust antiviral protection. However, pluripotent stem cells (PSCs) present a stark deviation from this canonical pathway. It has been shown that PSCs, including embryonic stem cells (ESCs) and induced pluripotent stem cells (iPSCs), do not produce IFNs in response to viral infection or poly(I:C) treatment and have a severely attenuated response to IFN treatment (5, 6). This resistance to IFN signaling is believed to be crucial for preserving the stem cells’ pluripotency and their ability to proliferate, as certain ISGs have been shown to have anti-proliferative and pro-apoptotic functions (3, 7, 8) and can induce differentiation (9, 10). Indeed, pluripotency and IFN signaling appear to be incompatible, as enforcing an IFN signaling in iPSCs disrupts their ability to differentiate equivalently into all three germ layers (11). Consistent with this, the pluripotency factors OCT4/POU5F1, SOX2, and KLF4 appear to be directly or indirectly involved in this repression (11).

Despite their lack of robust IFN responses, PSCs are not defenseless and are often described to be resistant to viral infection (10). In addition to reported vestigial RNA interference activity (12–14), PSCs rely on intrinsic expression of antiviral genes, including known ISGs, to confer antiviral resistance (15, 16). The mechanistic underpinnings of how PSCs balance the suppression of the IFN system while still intrinsically expressing immune-related genes remain largely unclear. Understanding this regulatory balance is crucial, as it allows stem cells to preserve their regenerative capacity while preventing viral takeover. As PSCs differentiate into somatic cells, they gradually acquire the ability to mount traditional IFN responses, indicating a dynamic regulation of IFN signaling during the differentiation process (15, 17).

In this study, we used single-cell RNA-sequencing (scRNA-seq) on virus-infected human ESCs (hESCs) to uncover the molecular regulators that govern antiviral and IFN responses in stem cells and investigate how these regulatory networks shift during differentiation to enable canonical IFN signaling. Our analyzes reveled a hESC subpopulation able to express IFN and ISGs upon Influenza A infection (IAV), suggesting a previously unrecognized heterogeneity in IFN response even within pluripotent populations. Our analyses revealed that negative regulators of the IFN response are intrinsically expressed in hESCs and under the control of pluripotency factors. Expression of these negative regulators is reduced in IFN-producing and responding cells. By elucidating these mechanisms, our work provides new insights into how stem cells navigate the delicate equilibrium between antiviral defense and their need to remain undifferentiated and proliferative.

## RESULTS

### Emergence of Inducible Immune Responses During Differentiation

To characterize how innate immune responses change during differentiation, we established two independent differentiation models (**Fig. 1A**). H1 human embryonic stem cells (hESCs) were differentiated toward lung progenitors, while H9 hESCs were differentiated toward cardiomyocytes (18, 19). For each lineage, we analyzed cells at defined intermediate stages of differentiation. For the lung progenitor lineage, we examined undifferentiated H1 cells (D0) as well as cells at Day 6 (D6, anterior foregut endoderm) and Day 15 (D15, lung progenitors) of differentiation. In parallel, for the cardiomyocyte lineage, we analyzed undifferentiated H9 cells (D0) together with cells collected at Day 5 (D5, cardiac mesoderm) and Day 13 (D13, cardiac progenitors) of differentiation. In both models, D0 represents the pluripotent state, whereas later time points capture progressively differentiated intermediate cell populations along each lineage trajectory. Cells at each differentiation stage were infected with an influenza A virus (H1N1/PR8) lacking the NS1 gene (IAVΔNS1) (20). Given that NS1 is IAV’s principal viral antagonist of host innate immunity, including IFN responses (21–25), IAVΔNS1 servs a potent stimulator of antiviral responses, enabling us to more clearly reveal differences in innate immune activation across pluripotent and differentiated cell stages.

**Fig. 1.**
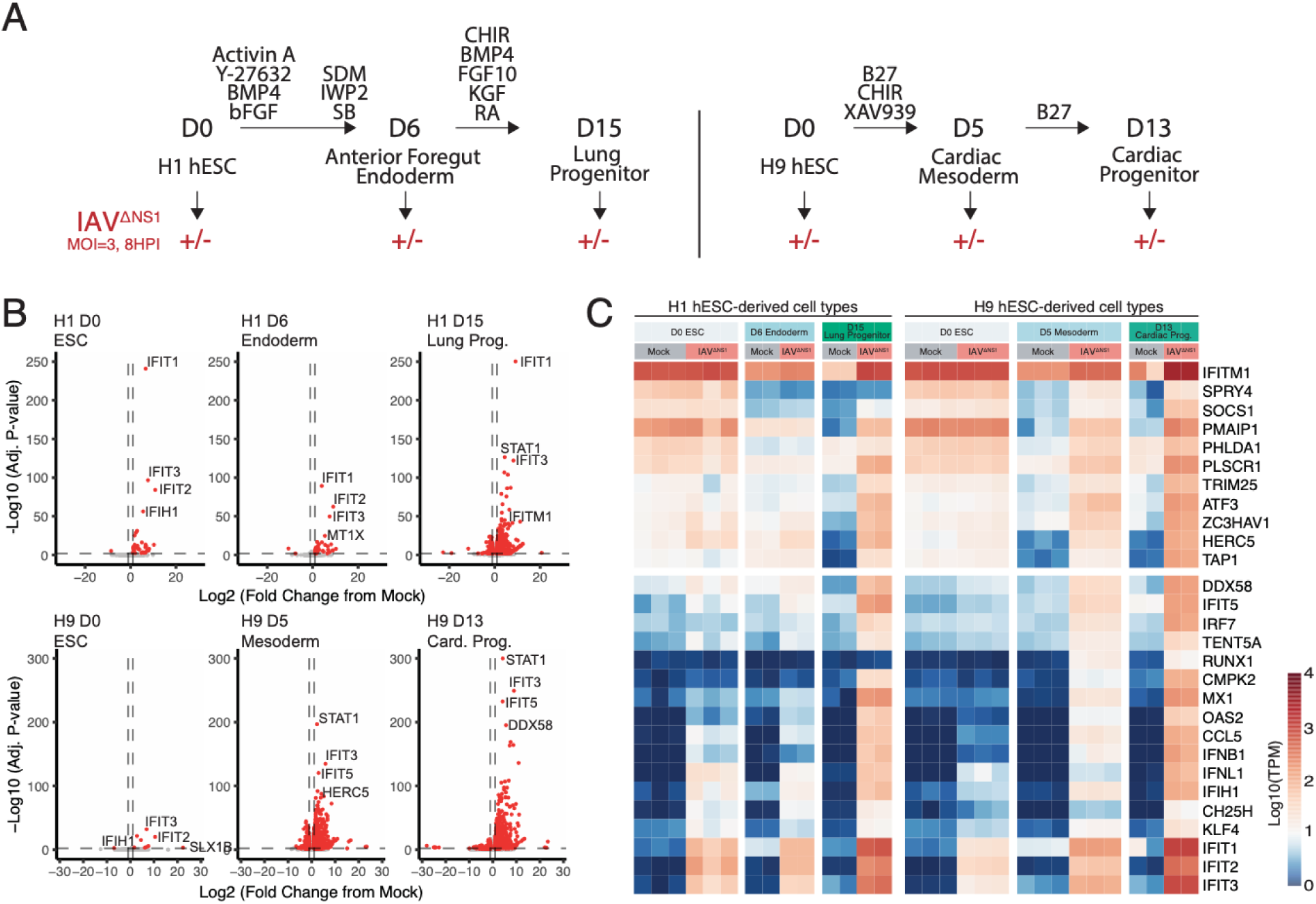
Immune response to virus infection progressively matures through differentiation. (**A**) Schematics summarizing the *in vitro* systems that model the hESC differentiation and its immune activation via IAVΔNS1 infection (18, 19). (**B**) Volcano plots indicating the transcriptional changes within each differentiation stage upon IAVΔNS1 infection, compared to the corresponding mock infection controls. The differentially expressed genes (|log_₂_ fold change| ≥ 1, adjusted *p* ≤ 0.05) are highlighted in red, with the top 4 up-regulated genes labeled in each plot. (**C**) Heatmap highlighting the expression levels of the key innate immune response genes, shown in each row. Each column represents a biological replicate, with its corresponding cell type and infection condition labeled at the top. Genes intrinsically expressed in ESCs are depicted in upper block. The color of the heatmap indicates the normalized read counts of each gene (log_10_ Transcripts per million).

Consistent with prior studies, the expression of most inflammatory genes remained largely unchanged relative to mock-infected controls in undifferentiated H1 and H9 hESCs (**Fig. 1B** and **Table S1 and S2**). Nonetheless, although hESCs are only moderately susceptible to infection with wild-type IAV (15), infection with IAVΔNS1 elicited subtle but reproducible transcriptional changes, with 34 and 9 genes upregulated in H1 and H9 D0 cells, respectivel (**Fig. 1B**). In contrast, differentiation toward lung and cardiomyocyte lineages progressively enhanced cellular responsiveness to viral infection, as evidenced by increasingly extensive transcriptional changes (**Fig. 1B** and **1C**). Intermediate differentiation stages exhibited 48 and 988 upregulated genes at D6 (H1) and D5 (H9), respectively, whereas the most differentiated populations mounted robust responses, with 1,351 and 1,133 upregulated genes at D15 (H1) and D13 (H9) (**Fig. 1B**). Consistent with this progression, well-characterized interferon-stimulated genes (e.g., IFIT1, IFIT2, IFIT3, MX1 and STAT1), along with PRRs DDX58/RIG-I and IFIH1/MDA5, were induced upon IAVΔNS1 infection; however, both the magnitude and statistical significance of their induction increased with advancing differentiation, reflecting the progressive maturation of the IFN response (**Fig. 1C**).

To assess whether this increased antiviral responsiveness during differentiation is the result of altered expression of core interferon signaling components, we analyzed the expression of key IFN pathway effectors across pluripotent and differentiated states. We examined the expression of PRRs relevant to IAVΔNS1 infection (DDX58/RIG-I, IFIH1/MDA5, MAVS), transcription factors involved in IFN production and inflammatory signaling (RELA, NFKB1, IRF3, IRF7), type I and III IFN receptors (IFNAR1, IFNAR2, IFNLR1, IL10RB), and downstream signal transducers (STAT1, STAT2, IRF9) (**Fig. S1A** and **S1B**). Surprisingly, comparison with D0 hESCs revealed only limited and inconsistent differences in the expression of these IFN pathway components across differentiation states. Notably, D15 lung progenitor cells exhibited lower expression of several effectors – including IFIH1, RELA, IFNAR2, IFNLR1, STAT2, and IRF9 – relative to their undifferentiated H1 counterparts (**Fig. S1A**), despite mounting a stronger antiviral response to IAVΔNS1 infection. Together, these observations indicate that the enhanced antiviral response observed upon differentiation is not driven by increased expression of IFN signaling components but instead suggest the existence of alternative regulatory mechanisms underlying the unique innate immune response in hESCs.

### Repetitive Element Expression Is Associated with Reduced Immune Sensing Capacity

PSCs are thought to restrain robust IFN-mediated immune responses in part because many classes of repetitive genomic elements – including long interspersed nuclear elements (LINEs), short interspersed nuclear elements (SINEs), and human endogenous retroviruses (HERVs) – are epigenetically derepressed in PSCs relative to differentiated somatic cells (10). These repetitive elements, which comprise a substantial fraction of the human genome and include retrotransposons of viral origin, can give rise to immunostimulatory RNAs and proteins when transcribed, and are normally kept silent by DNA methylation and chromatin-based repression to preserve genomic stability and prevent inappropriate innate immune activation (26, 27). Expression of such elements has been implicated in triggering innate immune pathways and cellular stress responses in other contexts, such as cancer (28, 29), and could therefore compromise pluripotency or trigger apoptosis if a full IFN response was engaged during the pluripotent state. To explore how this potential source of immune stimulation changes through differentiation, we analyzed the expression of repetitive elements in uninfected cells across our H1 and H9 differentiation lineages via bulk RNA-seq (**Fig. S2**). We found that upon differentiation there is a statistically significant reduction in HERV expression for both H1 and H9 lineages, when compared to D0 hESCs (**Fig. S2**). In fact, a statistically significant decrease in the expression of LINEs and SINEs is also detectable in H1 differentiation (**Fig. S2A**). These findings support a model in which IFN signaling is restrained in pluripotent stem cells to prevent inappropriate immune activation driven by endogenous repetitive element expression. However, despite this constraint, hESCs retain antiviral protection by intrinsically expressing a distinct set of immune-related genes.

### Intrinsic immunity in hESCs is regulated by pluripotency factors

In agreement with previous reports, we observed a set of ISGs (e.g., IFITM1, TRIM25, HERC5, ZC3HAV1/ZAP) that are intrinsically expressed in uninfected D0 hESCs and showed little to no induction following IAVΔNS1 infection (**Fig. 1C**). To fully characterize these immune-related genes that are intrinsically expressed in undifferentiated hESCs (D0), we expanded our analysis beyond ISGs to encompass genes associated with acquired immune responsiveness upon differentiation. We first compared transcriptomes of D0 hESCs with those of the most differentiated stages (i.e., D15 for H1 and D13 for H9) under mock conditions, thereby defining genes intrinsically expressed in undifferentiated cells but not in their differentiated counterparts (**Fig. 2A**, **Fig. S3A** and **Table S1 and S2**). To capture the immune potential of differentiated cells with intact IFN responsiveness, we identified genes induced upon viral infection by comparing mock– and IAVΔNS1-infected D15 (H1) and D13 (H9) populations (**Fig. 2A**, **Fig. S3A** and **Table S1 and S2**). The intersection of these datasets yielded 712 genes in H1 and 348 genes in H9 that are induced during a mature antiviral response yet intrinsically expressed in undifferentiated hESCs; we defined this set as intrinsic immune genes (**Table S3**). Notably, only a subset of these genes – 49 in H1 and 46 in H9 – corresponded to previously defined ISGs (30) (**Fig. 2B**, **Fig. S3B** and **Table S3**). The remaining intrinsic immune genes were enriched for processes related to transcriptional regulation, as well as pathways involved in cell cycle control and negative regulation of apoptosis or cell death (**Fig. 2C** and **Fig. S3C**). In H9 cells, intrinsic immune genes additionally showed enrichment for NF-κB signaling–related terms, suggesting lineage– or cell type-specific immune regulatory features (**Fig. S3C**). Consistent with a close link between intrinsic immunity and pluripotency, enrichment analysis of ChIP-seq datasets revealed that the pluripotency-associated transcription factors NANOG, SOX2, and POU5F1/OCT4 were consistently enriched regulators of intrinsic immune genes across multiple datasets in both H1 and H9 hESCs (**Fig. 2D** and **Fig. S3D**). Together, these results support a model in which intrinsic immune gene expression in hESCs is directly integrated into the pluripotency transcriptional network.

**Fig. 2.**
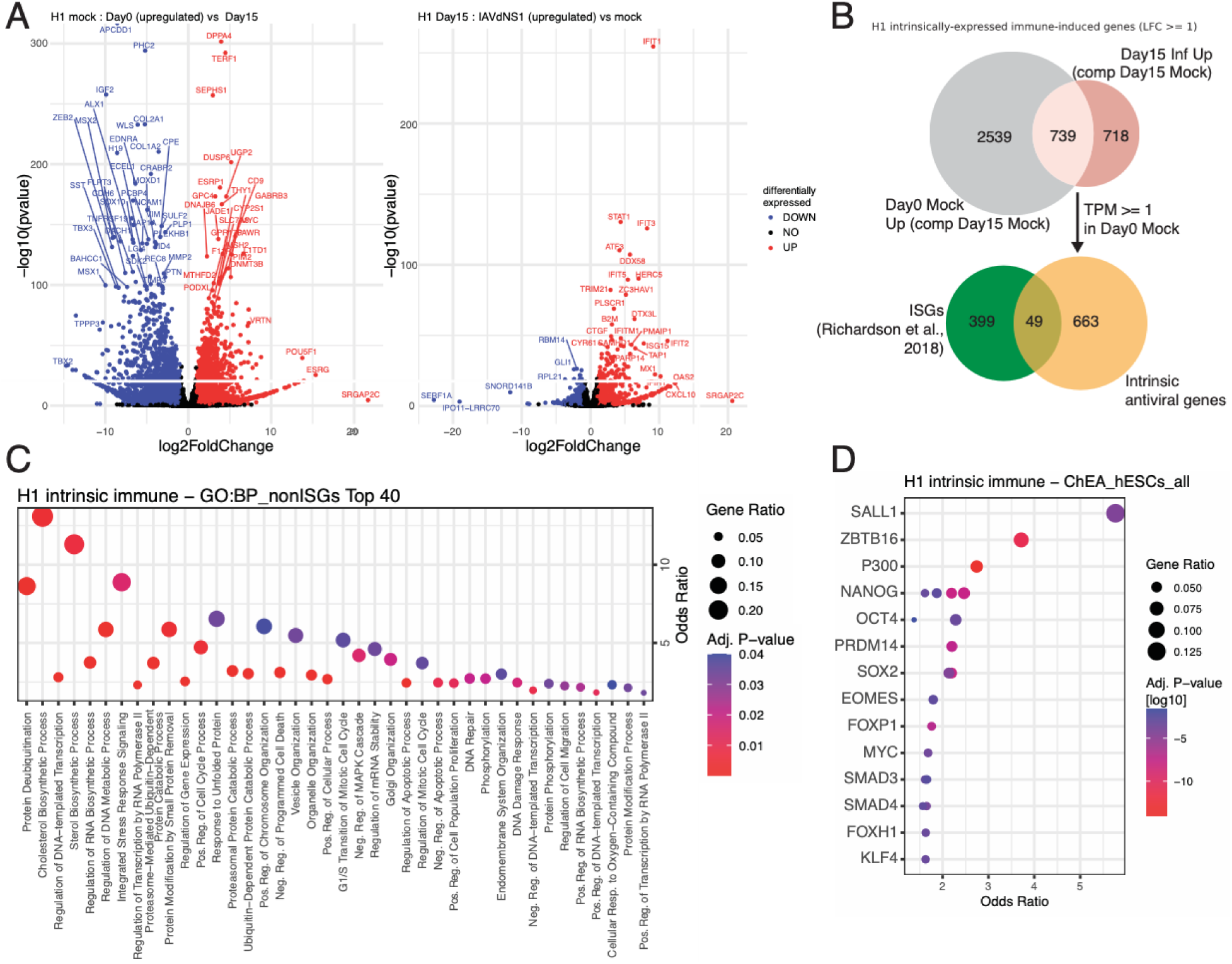
Defining intrinsic immune genes in H1 hESCs. (**A**) Volcano plots highlighting genes upregulated in D0 hESCs compared to D15 lung progenitors (left side) and genes upregulated in D15 lung progenitors by IAVΔNS1 infection (right side). Differentially expressed genes (|log_₂_ fold change| ≥ 1, adjusted *p* ≤ 0.05) are colored in blue (downregulated) and red (upregulated). (**B**) Venn diagram depicting the definition of intrinsically expressed immune genes. (**C**) Top 40 Gene ontology (GO) Biological Processes (BP) terms enriched in intrinsically expressed, non ISG immune genes. (**D**) ChEA 2022 enrichment results for ChIP-Seq experiments in hESCs for all intrinsic immune genes in H1 hESCs. Each dot corresponds to one ChIP-Seq experiment.

### Single-cell analysis uncovers a hESC subpopulation that exhibits IFN responsiveness

Bulk RNA-seq analyses revealed that hESCs mount a distinct and overall attenuated innate immune response to IAVΔNS1 infection. While the inducibility of most immune-related genes was strongly repressed, hESCs maintained high basal expression of intrinsic immune genes, consistent with a largely static antiviral state. Nevertheless, a small subset of immune-related genes – such as DDX58/RIG-I, MX1, OAS2, and IFIT1 – showed modest induction upon infection in D0 hESCs (**Fig. 1B** and **1C**). These observations suggest that, although bulk responses in pluripotent cells are minimal, some degree of inducible immune signaling is present. Given the potential heterogeneity within hESC populations, we next sought to resolve whether this limited induction reflects responses from a distinct subset of cells by examining gene expression at single-cell resolution.

We performed 3′ scRNA-seq on mock– and IAVΔNS1-infected H1 D0 and D6 cells, as the D6 anterior foregut endoderm stage represents a true intermediate between the attenuated immune responsiveness of undifferentiated hESCs and the robust immune sensing observed in D15 lung progenitors (**Fig. 1B**). Notably, D6 cells already exhibit reduced expression of intrinsic immune genes relative to D0 hESCs (**Fig. 1C**), indicating that maturation of the IFN response has initiated at this stage. Force-directed layout (FDL) representation of single-cell transcriptomes (**Fig. 3A** and **3B**) showed that D0 hESCs formed a largely homogeneous population characterized by high expression of core pluripotency markers (e.g., NANOG) as well as intrinsic immune genes (**Fig. 3C**, **Fig. S4A, S4B** and **S4C**). Expression of both pluripotency and intrinsic immune genes decreased upon differentiation, as observed in D6 mock-treated cells (**Fig. 3C**, **Fig. S4A** and **S4C**). The transcriptional homogeneity of mock D0 hESCs further indicates minimal spontaneous differentiation within the pluripotent cultures. The FDL also revealed that differentiation resulted in three transcriptionally distinct populations at the D6 stage (**Fig. 3A** and **3B**). Examination of cell type–specific marker expression under mock conditions within the D6 population identified cell subpopulations expressing endoderm lineage markers, such as PRTG (31), alongside more committed progenitors expressing lung-associated markers, including FOXA2 (32), highlighting the emergence of lineage commitment at this stage (**Fig. 3C**).

**Fig. 3.**
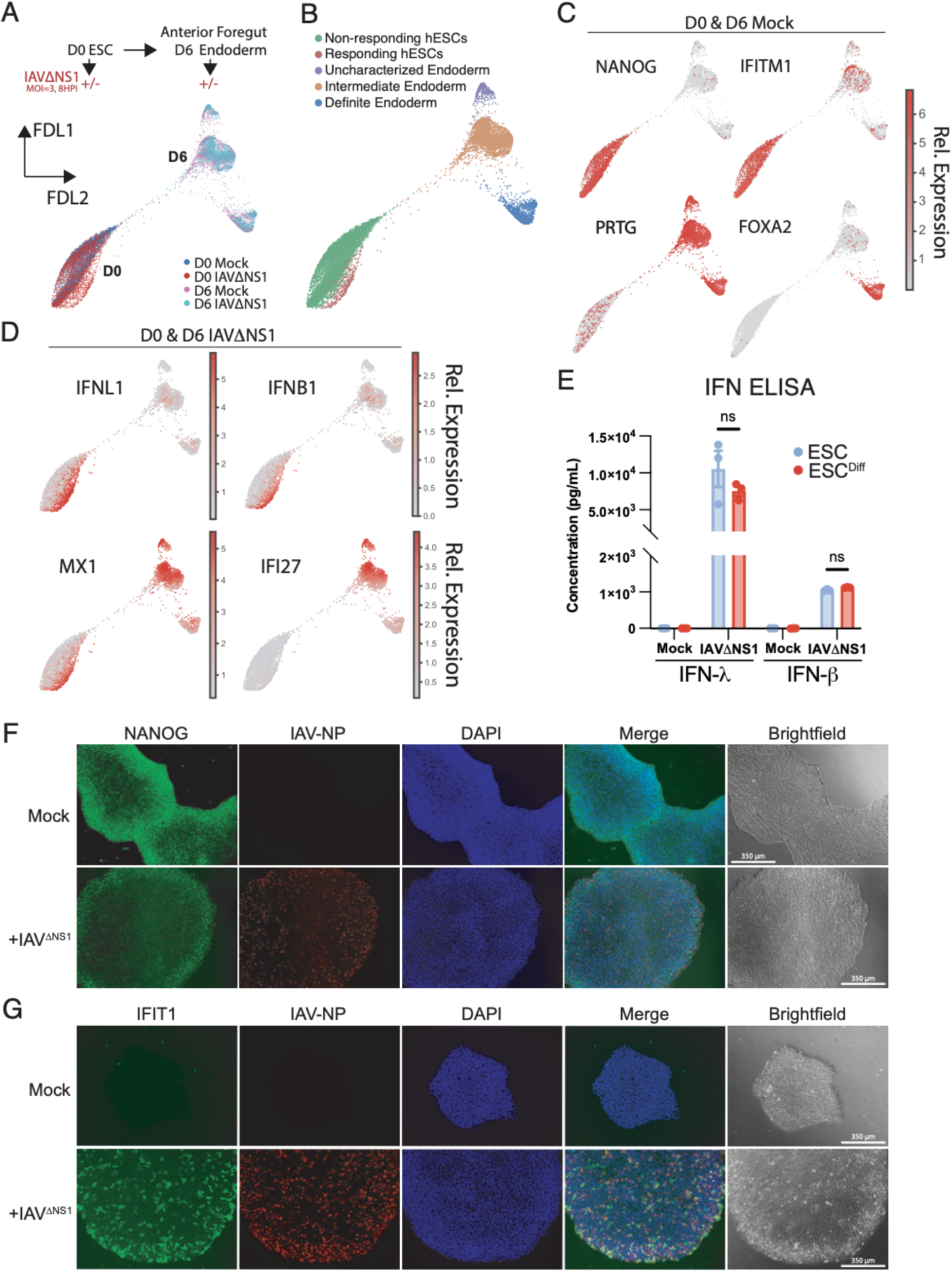
Single cell transcriptomic analysis reveals a subset of IFN-responsive hESCs upon viral infection. (**A**) Diagram and force-directed layout (FDL) summarizing the experimental setup of the scRNA-seq experiment. D0 H1 hESC and differentiated D6 anterior foregut endoderm cells were independently infected with IAVΔNS1 at MOI of 3 for 8 hours. (**B**) Cell annotation according to Leiden clusters based on their transcriptional proximity. (**C**) Expression of cell marker genes and an intrinsic immune gene (IFITM1) in mock cells highlight the effect of differentiation resulting in distinct cell populations. Relative expression indicates MAGIC-imputed, log-normalized read counts. (**D**) Expression of IFN genes and ISGs in IAVΔNS1 infected cells. (**E**) ELISA quantification of secreted IFNλ and IFNβ in the supernatant of IAVΔNS1 infected undifferentiated H1 cells (ESC) or after FBS-induced differentiation (ESC^Diff^). Supernatant was harvested at 16hpi. (**F**) Immunofluorescence assay depicting expression of NANOG and viral protein NP in mock and IAVΔNS1 infected D0 hESC colonies. (**G**) Immunofluorescence assay depicting expression of IFIT1 and viral protein NP in mock and IAVΔNS1 infected D0 hESC colonies. Scale bar = 350 μm.

To assess innate immune activation at single-cell resolution, we examined the expression of key IFN response genes in IAVΔNS1-infected cells, including type I and III IFNs (IFNB1 and IFNL1) and representative ISGs (**Fig. 3D** and **Fig. S5**). Unexpectedly, we identified a distinct subset of infected D0 hESCs – absent in mock-treated controls – that expressed both IFNs and ISGs (**Fig. 3B**, **3D** and **Fig. S5A**). In D6 cells, IFN expression was confined to a small subset of cells but was associated with widespread ISG induction across cell populations, consistent with effective paracrine signaling (**Fig. 3D** and **Fig. S5**). In contrast, this paracrine amplification was not observed in D0 hESCs (**Fig. 3D** and **Fig. S5**). While D0 hESCs contained a larger fraction of IFN-expressing cells, ISG induction was confined almost exclusively to these IFN-expressing cells, suggesting that paracrine IFN signaling fails to propagate efficiently in hESCs. To verify the expression of IFN proteins, we performed ELISA to measure IFNλ1/3 and IFNβ levels in the supernatant of IAVΔNS1-infected H1 hESCs (**Fig. 3E**). Remarkably, D0 hESCs produced levels of IFN protein comparable to those observed in IFN-responsive differentiated cells. Importantly, we did not observe any correlation between the virus burden (measured in viral UMI counts) and the responsiveness state of D0 hESCs (**Fig. S5B** and **S5C**).

To assess whether hESCs possess the core components required for IFN signaling, we examined the expression of key elements in this pathway between responding and non-responding hESCs (**Fig. 3B**), including IFN receptors (IFNAR1, IFNAR2, IFNLR1, and IL10RB), signal transducers (STAT1, STAT2, JAK1, TYK2), and pattern recognition receptors (DDX58/RIG-I, IFIH1/MDA5, MAVS, and TLR3) (**Fig. S4D** and **S4E**). While expression of most components was comparable between responding and non-responding hESCs, several key factors (i.e., STAT1, DDX58/RIG-I, and IFIH1/MDA5) were strongly upregulated in responding cells upon infection. Importantly, these factors were already detectable in non-responding hESCs, indicating that the limited IFN response cannot be attributed to their absence but rather to restricted signaling.

To exclude the possibility that IFN signals originated from residual spontaneously differentiated cells, we performed immunofluorescence analysis to confirm that hESC colonies uniformly expressed high levels of the pluripotency marker NANOG (**Fig. 3F**). Following IAVΔNS1 infection, viral nucleoprotein (NP) staining revealed comparable infection of cells at both the center and periphery of hESC colonies (**Fig. 3F**). Notably, a subset of infected cells across the colony exhibited high IFIT1 expression, demonstrating that the ISG signal observed by scRNA-seq did not arise from spontaneously differentiated cells (**Fig. 3G**). Importantly, nearly all IFIT1-positive cells were also positive for viral NP, indicating that active viral replication is required to trigger immune activation in hESCs (**Fig. 3G** and **Fig. S6A).** To confirm the specificity of NANOG immunostaining, we deliberately retained spontaneously differentiating cells in hESC cultures and observed a complete loss of NANOG signal in these cells, verifying that NANOG staining reliably marks pluripotent cells (**Fig. S6B**). Under these conditions, only spontaneously differentiated cells – but not pluripotent hESCs – responded to IFNβ treatment (i.e., showed IFIT1 induction) (**Fig. S6C**), in agreement with the block of paracrine signaling in hESCs. These results support our single-cell observations that inducible IFN responses in hESC cultures are confined to a subset of infected cells and do not arise from contaminating differentiated populations.

Finally, we found that this transcriptional response was not limited to IAVΔNS1 infection, as Sendai virus (SeV) elicited a comparable induction of IFNB, IFNL1, IFIT1, and MX1 while pluripotency markers (i.e., NANOG, SOX2, and POU5F1/OCT4) remained at baseline levels (**Fig. S7A**). In contrast, infection with wild-type IAV failed to elicit a detectable response, suggesting that the immune activation observed depends on the absence, or functional impairment, of viral IFN antagonists such as NS1. Notably, the magnitude of gene induction did not correlate with the abundance of viral RNA, as levels of IAV nucleic acids (IAV N) were more than 60-fold higher than those of SeV (SeV NP), despite eliciting comparable transcriptional responses (**Fig. S7A** and **S7B**).

In summary, single-cell RNA-seq analyses indicate that H1 D0 hESCs are capable of producing IFNs upon virus infection. ISG induction is confined to this same subset of IFN-expressing cells, whereas non-responsive hESCs remain transcriptionally indistinguishable from uninfected controls (**Fig. 3A**, **3B**, and **3D**). Importantly, while we cannot fully attribute the observed transcriptional program to autocrine IFN signaling – since several targets, including IFIT1, DDX58/RIG-I and IFIH1/MDA5, can be induced directly downstream of IRF3/IRF7 activation following viral sensing, independent of JAK–STAT signaling (**Fig. S5A**) (4, 33) – the robust expression of canonical IFN-signaling dependent genes such as MX1 and IFI6, which are associated with ISGF3-mediated transcription downstream of IFN receptor engagement (**Fig. 3D** and **S5A**) (34, 35), supports a contribution of autocrine IFN signaling within this responsive subset. In contrast to bulk RNA-seq analyses, single-cell profiling reveals a pronounced bimodality in innate immune activation within hESC cultures. Because most cells do not exhibit transcriptional responses, this heterogeneity is obscured at the bulk level and manifests as an overall attenuated immune response (**Fig. 1**). Identification of this immune-responsive subpopulation enables further interrogation of the regulatory mechanisms that constrain IFN signaling directly in undifferentiated D0 hESCs.

### Attenuation of IFN Negative Regulators Enables Paracrine IFN Signaling in hESCs

While immunofluorescence analyses and pluripotency gene expression scores indicate that responding hESCs largely retain pluripotent identity (**Fig. 3F** and **S4C**), we nonetheless observed subtle remodeling of the pluripotency program. Specifically, expression of the pluripotency factor SOX2 was reduced in responding hESCs relative to non-responding cells, whereas NANOG and OCT4/POU5F1 expression remained largely stable (**Fig. 4A**). To systematically characterize transcriptional changes associated with the transition to the IFN-responsive phenotype, we used trajectory inference on single cell transcriptomes to map cell to cell relationships in D0 IAVΔNS1-infected hESCs. Cells were ordered along a pseudotime continuum spanning from a pluripotent state, defined by high NANOG expression, to an immune-responsive state, defined by elevated expression of genes enriched in responding hESCs (**Fig. 4B**). Analysis of gene expression dynamics along this trajectory revealed distinct transcriptional programs associated with progression toward the responsive state (**Fig. 4C**, **4D** and **Table S4**). Genes showing the strongest induction at late pseudotime (i.e., responsive state) formed a cluster enriched for antiviral and IFN-related genes, including IFNL1, IFIT1, and MX1 (**Fig. 4C**, **4D** and **Table S4**). In contrast, pluripotency factors followed divergent but uniformly declining expression trends, with NANOG (cluster 0), SOX2 (cluster 4), and POU5F1/OCT4 (cluster 6) occupying distinct clusters that all decreased as cells approached the immune-responsive state (**Fig. 4C** and **Table S4**). Notably, the cluster containing SOX2 was enriched for genes associated with Wnt signaling (**Fig. 4C** and **4D**), a pathway with established roles in regulating pluripotency across developmental contexts (36). Together, these analyses suggest that acquisition of immune responsiveness in hESCs is accompanied by coordinated, yet modest, remodeling of the pluripotency transcriptional program.

**Figure 4.**
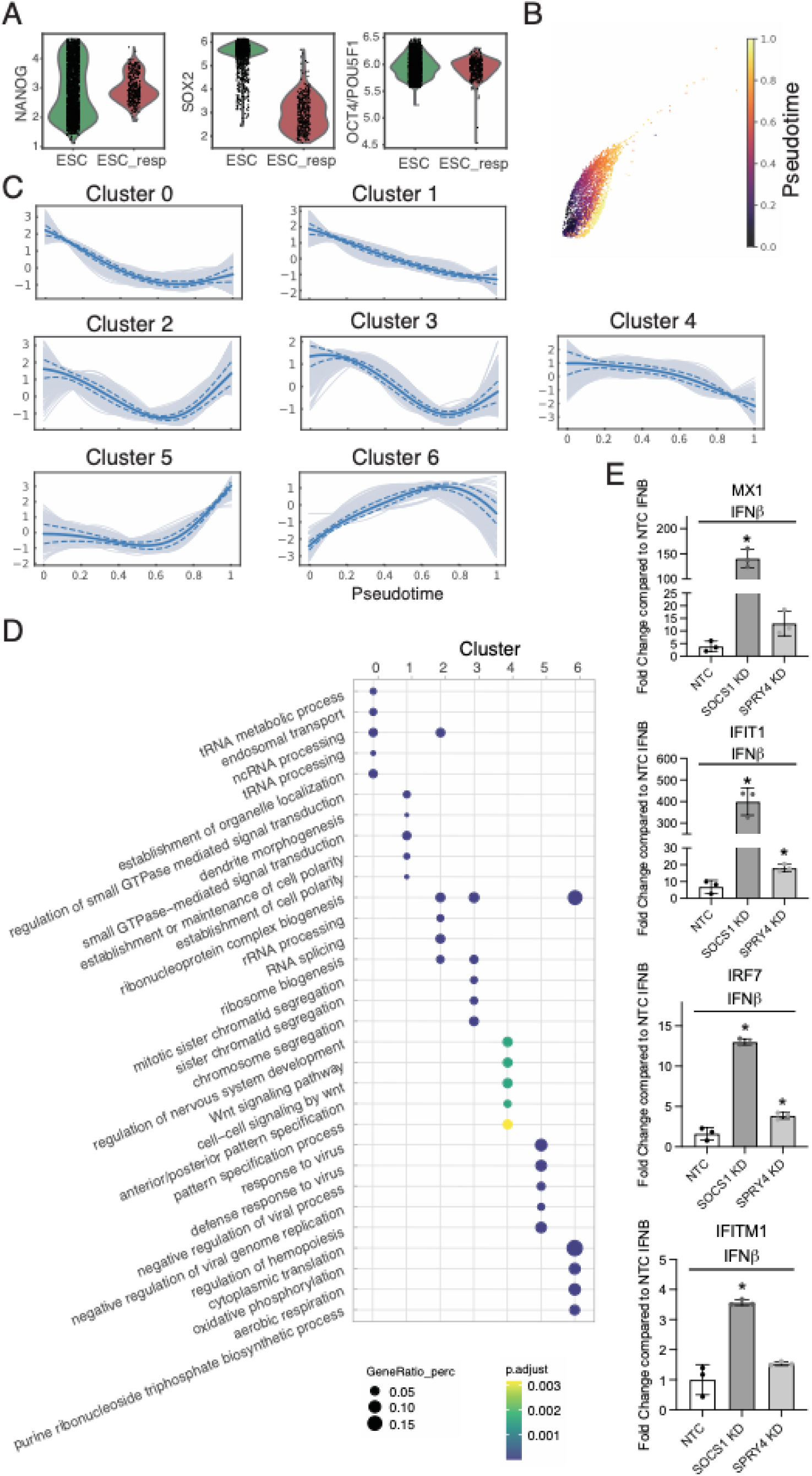
Comparative analysis between responder and non-responder populations. (**A**) Expression of pluripotency factors in D0 IAVΔNS1 infected H1 hESCs. Responsive and non-responsive hESCs as defined in Fig. 3B, respectively. (**B**) Pseudotime analysis of D0 IAVΔNS1-infected ESCs. The pseudotime trajectory was anchored by defining the start cell as the cell with the highest NANOG expression and the terminal cell as the cell exhibiting the highest expression score for the top 200 genes differentially expressed in responsive hESCs. (**C**) Clustering of gene expression trends along pseudotime. Both pseudotime analysis and gene trend dynamics were generated using Palantir (40). (**D**) Gene Ontology Biological Processes (GO BP) enriched for each cluster from (C). (**E**) RT-qPCR quantification of MX1 and IFIT1 after siRNA-mediated knock-down (KD) of SOCS1 and SPRY4 and IFNβ treatment (100 U/ml, 8hpt). Asterisks indicate p-value < 0.05 compared to NTC control with Welch’s unpaired t-test.

Notably, in addition to pluripotency factors such as NANOG and SOX2, gene clusters that decline along pseudotime (clusters 0 and 4) also contained multiple factors known to restrain interferon signaling (37), including SOCS1, SPRY1, and SPRY4 (**Fig. S8A** and **Table S3**). This synchronous downregulation raised the possibility that relief of intrinsic inhibitory mechanisms contributes to the emergence of responsive phenotypes. To test this hypothesis, we independently knocked down SOCS1 and SPRY4 using siRNAs and assessed the ability of hESCs to respond to paracrine IFN signaling following IFNβ treatment (**Fig. 4E** and **Fig. S8B**). Quantification of immune activation by RT-qPCR revealed robust induction of ISGs (i.e., IFIT1, MX1, IRF7, and IFITM1) upon SOCS1 knockdown compared to non-targeting controls (**Fig. 4E**). Knockdown of SPRY4 also enhanced expression of select ISGs, such as IFIT1 and IRF7, albeit to a lesser extent than SOCS1 depletion (**Fig. 4E**). These differences are consistent with the distinct mechanisms by which these regulators constrain IFN signaling. SOCS1 inhibits JAK-mediated STAT1 phosphorylation, a central step in canonical ISG induction (38), whereas SPRY4 has been implicated in limiting p38 MAPK–dependent recruitment of CBP/p300 to ISG promoters (39). Importantly, neither knockdown nor IFNβ treatment altered expression of core pluripotency factors (**Fig. S8C**).

Remarkably, SOCS1 was identified as an intrinsic immune gene in both H1 and H9 hESCs, while SPRY4 were intrinsic immune genes in H9 hESCs (**Table S3**). Consistent with the regulation though pluripotency factors observed for other intrinsic immune genes (**Fig. 2D**), analysis of ChIP-seq datasets revealed binding of the pluripotency factor NANOG in promoter or enhancer regions proximal to SOCS1 and SPRY4 (**Fig. S9**). Together, these results indicate that key negative regulators of IFN signaling are directly embedded within the pluripotency transcriptional network, providing a mechanistic link between maintenance of pluripotency and suppression of paracrine interferon responses.

## DISCUSSION

In this study, we define how innate immune responsiveness emerges during differentiation of hESCs and identify transcriptional mechanisms that actively restrain IFN signaling in the pluripotent state. Using two hESC lines (H1 and H9) differentiated along lung and cardiac lineages, we show that pluripotent cells exhibit a markedly attenuated antiviral response compared with differentiated progeny. Infection with IAVΔNS1 – a potent activator of host antiviral signaling – elicited minimal transcriptional changes in undifferentiated hESCs, whereas progressively differentiated cells mounted increasingly robust immune responses, culminating in widespread induction of immune genes in lung and cardiac progenitors (**Fig 1B**). These findings reinforce the concept that acquisition of innate immune competence is a differentiation-dependent process tightly coupled with the exit from pluripotency.

A key insight from our single-cell analyses is that innate immune activation in hESCs is bimodal. While the majority of non-responding infected hESCs remained transcriptionally indistinguishable from uninfected controls, a distinct subpopulation of undifferentiated hESCs expressed IFN genes upon infection (**Fig. 3D**). Notably, the induction of ISGs was restricted to this same subset of IFN-expressing cells (**Fig. 3D** and **Fig. S5A**). This immune behavior, obscured in bulk analyses, reveals that inducible immune activation in hESCs is highly constrained and fails to propagate broadly through paracrine signaling, in stark contrast to differentiated cells. Importantly, these data suggest that IFN production alone does not drive downstream responsiveness; rather, only a distinct subpopulation possesses the competence to both produce IFN and execute the broader immune transcriptional program.

Mechanistically, we show that immune restriction in hESCs results from active repression rather than a lack of core signaling components. Trajectory analysis revealed that emergence of immune responsiveness is accompanied by subtle attenuation of the pluripotency transcriptional program and concurrent downregulation of negative regulators of IFN signaling, including SOCS1 and SPRY4 (**Fig. 4A** and **4C**). Functional perturbation experiments demonstrate that depletion of these factors – particularly SOCS1 – restores the capacity of hESCs to respond to paracrine IFN stimulation (**Fig. 4E**). These results are consistent with previous reports that implicated SOCS1 in the restriction of IFN signaling in iPSCs and mesenchymal stem cells ESCs (6, 41, 42). Our analyses further indicate that SOCS1 and SPRY4 are not only intrinsically expressed in hESCs together with other intrinsic immune genes but are also directly embedded within the pluripotency transcriptional network and regulated by the key pluripotency factor NANOG (**Fig 2B**, **Table S3** and **Fig. S9**). These findings provide a mechanistic framework linking maintenance of pluripotency and active repression of IFN signaling, enabling pluripotent cells to balance antiviral protection with preservation of developmental potential.

While our analyses focused on two differentiation trajectories and hESC lines, immune regulation may differ across additional lineages or in iPSC systems, and further studies are needed to generalize these findings. Nonetheless, our work advances our understanding of how IFN signaling is selectively constrained in pluripotent cells and reveals that negative regulators of IFN responses are integral components of the pluripotency program itself. At the same time, our findings raise important open questions. First, it remains unclear what precisely drives individual pluripotent cells to transition from a fully pluripotent state into one that permits pathogen sensing, IFN production, and immune gene induction. One possibility is that exceptionally strong pathogen-associated signals are required to overcome the restrained sensing capacity of hESCs. Consistent with this idea, both IAVΔNS1 and SeV generate abundant defective viral genomes (DVGs) during infection, which are potent ligands for RIG-I–like receptors and other innate sensors (43). Accumulation of such defective RNAs, in combination with impairment of viral antagonists (i.e., IAV NS1) (44), may therefore saturate otherwise constrained pathogen recognition pathways and enable activation of antiviral programs in a small subset of cells (**Fig. S7**). Second, previous work has shown that enforced engagement of IFN signaling in PSCs impairs their differentiation potential (11), yet the ultimate fate of these immune-responsive hESCs remains unclear – whether these cells undergo cell death, experience lasting differentiation defects, or adopt alternative developmental trajectories. Third, the existence of a rare immune-responsive subpopulation raises the intriguing possibility that such cells may serve a protective role within the pluripotent population, potentially engaging strong antiviral programs at the cost of their own developmental fitness to safeguard surrounding cells in the early embryonic microenvironment. Addressing these questions will be critical for understanding how innate immunity and developmental potential are balanced during early human development. Beyond stem cell biology, these insights have broader implications for IFN regulation, highlighting strategies by which antiviral defenses can be uncoupled from excessive inflammatory signaling. Importantly, defining mechanisms that transiently relieve immune repression without disrupting pluripotency may inform the development of safer and more effective stem cell-based therapies.

## MATERIALS AND METHODS

### Cell culture, treatment and viral infection

Human ESC line H1 (WA01, H1 hESC) was obtained from WiCell Research Institute, Inc.. All H1 hESC lines are maintained in Vitronectin XF (STEMCELL 07180)-coated plates in mTeSR Plus media (STEMCELL 100-0276), with routine media change every other day. All cells were maintained at 37°C and 5% CO2. All subcultures of hESC were passaged using Gentle Cell Dissociation Reagent (STEMCELL 100-0485). CloneR2 was added to hESCs directly after passaging for 24hours. For single-cell analyses, Human ESC lines H1 and H9 (WiCell Research Institute WA09) were maintained on 1% Matrigel (Corning)-coated six-well plates in StemFlex medium (Gibco) at 37 °C with 5% CO_2_, as previously described (19).

Influenza A/Puerto Rico/8/34 (H1N1) viruses with NS1 deletion (20) (IAVΔNS1, NCBI:txid183764) were grown for 48 h in MDCK cells (for IAV-WT) or in MDCK-NS1-GFP cells (for IAVΔNS1) (45) at an MOI of 0.05 in DMEM supplemented with 0.3% BSA (MP Biomedicals®) and 1 μg/mL TPCK-trypsin (Sigma-Aldrich®). The infectious titer of the virus stock was determined using plaque assays in MDCK or MDCK-NS1-GFP cells. For infections using Sendai Virus (SeV) we used the Cantell strain (ATCC VR-907) and titers were calculated by hemagglutinin units (HAU). Infection of hESC and hESC-derived cells was performed at the indicated MOIs for 1 hour at 37°C in mTeSR Plus media before being replaced with fresh media and incubated for the indicated hours post infection at 37°C.

### Directed differentiation of H1 and H9 hESCs

Monolayer-based cardiomyocyte (CM) differentiation of H9 hESCs was perfomed as previously described (18). In brief, H9 hESCs were passaged at density of 3×10^5^ cells/well on a 6-well plate and grown for 48 h to 90% confluence. On day 0, the medium was replaced with RPMI 1640 supplemented with B27 without insulin and 6 μM CHIR99021. On day 2, the medium was changed to RPMI 1640 supplemented with B27 without insulin for 24 h. Day 3, medium was refreshed to RPMI 1640 supplemented with B27 without insulin and 5 μM XAV939 for another 48 h. On day 5, the medium was changed back to RPMI-B27 without insulin for 48 h, and then switched to RPMI 1640 plus normal B27 for another 48 h. On day 9, the medium was transiently changed to RPMI 1640-B27 without D-glucose for two days to allow metabolic purification of CMs. From that day on, fresh RPMI 1640-B27 was changed every two days. Mock and IAVΔNS1 infections were performed on D0, D5, and D13 cells before their corresponding media changes.

H1 hESCs differentiation into lung progenitors was performed as previously described (19). In brief, H1 hESCs were treated with Accutase and plated onto low-attachment 6-well plates (Corning), and then resuspended in endoderm induction medium – composed of serum-free differentiation (SFD) medium of DMEM/F12 (3:1) (Life Technologies) supplemented with 1x N2 (Life Technologies), 1x B27 (Life Technologies), 50 μg/ml ascorbic acid, 2 mM Glutamax (Gibco), 0.4 μM monothioglycerol, 0.05% BSA, 10 μM Y-27632, 0.5 ng/ml human BMP4 (R&D Systems), 2.5 ng/ml human bFGF and 100 ng/ml human activin A (R&D Systems) – for 72-76 h dependent on the formation rates of endoderm cells. On day 3 or 3.5, the endoderm bodies were dissociated into single cells using 0.05% trypsin/0.02% EDTA and plated onto fibronectin-coated, 24-well tissue culture plates (∼100,000–150,000 cells per well). For induction of anterior foregut endoderm (D6), the endoderm cells were cultured in SFD medium supplemented with 1.5 μM dorsomorphin dihydrochloride (R&D Systems) and 10 μM SB431542 (R&D Systems) for 36 h and then switched for 36 h to 10 μM SB431542 and 1 μM IWP2 (R&D Systems) treatment. For induction of early-stage lung progenitor cells (D15), the resulting anterior foregut endoderm was treated with 3 μM CHIR99021 (CHIR, Stem-RD), 10 ng ml^−1^ human FGF10, 10 ng ml^−1^ human KGF, 10 ng ml^−1^ human BMP4 and 50–60 nM all-trans retinoic acid (ATRA), in SFD medium on day 8-10. The day 10-15 culture was maintained in a 5% CO_2_/air environment. Mock and IAVΔNS1 infections were performed on D0, D6, and D15 cells before their corresponding media changes.

### Bulk RNA-seq

Infected and mock-infected cells were lysed in TRIzol (Invitrogen), following total RNA extraction with DNase I treatment using a Direct-zol RNA miniprep kit (Zymo Research) according to the manufacturer’s instructions. RNA-seq libraries of polyadenylated RNA were prepared using the TruSeq RNA Library Prep Kit v2 (Illumina) according to the manufacturer’s instructions and sequenced on an Illumina NextSeq 500 platform. Raw sequencing reads were trimmed for quality and adapters using bbduk (BBMap v39.03) with parameters ktrim=r qtrim=10 k=21 mink=11 hdist=2 maq=10 minlen=25 tpe tbo. Salmon (v1.9.0) (46) was used to quantify transcript expression using the trimmed reads against the human reference transcriptome (NCBI RefSeq hg38) with default parameters. Transcript-level abundance estimates were imported into R and summarized to gene-level counts using tximport (47). Differential gene expression analyses were then performed on these gene-level count matrices using DESeq2 (v1.44.0) (48). For the analysis of repetitive elements, raw sequencing reads were concatenated and processed using a custom nextflow pipeline, which performs both STAR alignment (v2.7) (49) and telescope (50) quantification of transposable elements (reassign_mode “choose”) against the human genome (hg38) and TEtranscripts (51) repetitive element dataset based on RepBase references (52). Counts for each element were summarized to the repeat class level and relative abundance (normalized by counts per million in edgeR (53)) was assessed in RStudio. Visualizations were generated using ggplot2.

### Single cell transcriptomics

Mock and IAVΔNS1-infected hESC and hESC-derived cells were treated with Accutase and collected as a single-cell suspension. Cells were then washed twice with ice-cold 1x PBS and filtered using a 40 μm Flowmi cell strainer (Bel-Art Scienceware). Cell count and viability were determined using trypan blue stain and a Countess II automatic cell counter (Thermo Fisher Scientific). Based on this cell count, a target cell input volume of 8,000 cells (Targeted cell recovery: 5,000 cells) were loaded into a Chromium Controller using Chromium NextGEM Single Cell 3′ v3.1 kit (10X Genomics) according to the manufacturer’s instructions. Library preparation proceeded after GEMs were generated following manufacturer’s instructions. Final libraries were then sequenced on an Illumina NovaSeq 6000. Sequencing data was processed with CellRanger v4.0.0 (10X Genomics). Reads were mapped to a combined human (GRCh38) and IAV PR8 (GenBank: AF389115.1, AF389116.1, AF389117.1, AF389118.1, AF389119.1, AF389120.1, AF389121.1, and AF389122.1) genome reference using CellRanger count. CellRanger output files were further analyzed with Scanpy. Standard quality control metrics were calculated using Scanpy’s *scanpy.pp.calculate_qc_metrics*. First, cells were filtered according to count numbers: 99.5th percentile as upper threshold and 37th, 56th, 64th and 74th percentile for lower threshold for D0 mock, D0 IAV ΔNS1, D6 mock and D6 IAV ΔNS1, respectively. Cells whose mitochondrial reads make up more than 20% of the total reads were filtered out. All samples were concatenated into one data object before further processing. Genes that are expressed in less than 50 cells, along with ribosomal and Influenza viral genes were filtered out. The expression data matrix was then log-normalized, and dimensionality reduced using principal component analysis (PCA). Force-directed layout (FDL) was generated from the PCA coordinates with Palantir, followed by imputation by MAGIC (40). Clustering was performed on the FDL using the Leiden algorithm. Each cell cluster is then annotated based on immune response (for D0 cells) or cell type-specific (for D6 cells) gene expression signatures. A pluripotency score was calculated for each cell based on gene expression of the following marker genes: NANOG, SOX2, THY1, DUSP6, LNCPRESS1, NTS, ZSCAN10, RFLNB, ADM, TRIML2, RAB17, GNG4, CD3EAP, SCG3, VASH2, ETV1, DUSP5, CYTOR and RRS1. An intrinsic antiviral gene score was computed for each cell based on the gene list from **Table S3**. **Palantir pseudotime analysis**

For computation of pseudotime, the preprocessed object was subset for cells from Leiden cluster 0 & 1 (non-responding hESCs) and 6 (responding hESCs) within D0 IAV ΔNS1 cells. A *responder* score was calculated based on the top 200 differentially expressed genes in the responding cluster compared to the non-responding hESCs cluster. For palantir pseudotime analysis, the cell with highest NANOG expression was used as starting cell, while the cell with the highest responder score was used as terminal cell. Taken together, palantir pseudotime shows the trajectory from highest NANOG expressor to most responding cell. Gene expression from all cells along the pseudotime trajectory was summarized as gene trend. Gene expression clusters were calculated for all genes that showed a variance greater than 0.01, with n_neighbors=150 and resolution=0.2.

### Enrichment analysis

Enrichment analysis of Gene Ontology Biological Processes (GO BP) terms and ChEA was performed with the Mayan Labs Enrichr software (54). For GO terms the GO_Biological_Process_2025 library was used and for ChEA the ChEA_2022 library (55). Enrichment was performed with genetic background (all genes expressed in the corresponding experiment).

### ELISA

ELISA was performed on cell culture supernatants obtained from IAVΔNS1-infected cells after 18 hours post infection to detect IFNβ (biotechne, #DY814-05) and IFNλ 1/3 (biotechne, #DY1598B-05). Protocol was performed according to the manufacturer’s instructions.

### Knock-down using siRNA

For efficient siRNA-mediated knock-down in hESCs, we double-transfected the cells as previously reported (56). 40 pmol of siRNA was diluted in 50 µl OPTIMEM medium and incubated for 5 min at room temperature. 5 uL Lipofectamine RNAiMAX (Thermo Fisher Scientific) was diluted in 50 uL OptiMEM and incubated for 5 min at room temperature. The diluted siRNA and Lipofectamine were mixed together and incubated for 30 min at room temperature. 100 µl of the combined mixture was added to 1×10^6^ freshly seeded H1 hESCs in a well of a 6 well plate. After 24 hours of incubation in a cell culture incubator at 37C, 5% CO2 and max humidity, medium was exchanged to fresh mTeSR+ medium. 48 hours after transfection, the process was repeated. Cells were further treated and/or harvested for RNA purification on the next day.

### Immunofluorescence

Antibodies and other reagents used for immunofluorescence can be taken from **Table 1**. hESCs were washed three times with PBS and then permeabilized and fixed with 90% ice-cold Methanol for 5 min. Cells were washed again with PBS, then incubated in blocking buffer (1 % BSA and 0.1 % Tween20 in PBS) for 1 hour at room temperature. Primary antibody was diluted in blocking buffer at 1:500 and incubated overnight at 4C. Cells were washed three times with blocking buffer before incubation with the secondary antibody in blocking buffer at 1:250 dilution. Incubation was performed at room temperature for 1 hour. Before imaging, cells were incubated in a 1:100 dilution of DAPI stain for 5 minutes and then washed three times in blocking buffer. Imaging was performed on Thermo Fisher EVOS M5000.

**Table 1.** Antibodies for immunofluorescence. Information for antibodies used in Fig. 3 and S6

| Antibody Target | Source | Manufacturer | Catalog Number |
| --- | --- | --- | --- |
| Nanog | Rabbit | Cell signaling | 3580S |
| Ifit1 (D2X9Z) | Rabbit | Cell signaling | 14769S |
| Influenza A virus, NP | Mouse | MilliporeSigma | MABF216425 |
| anti-Mouse AF594 | Donkey | Invitrogen | R37115 |
| Anti-Rabbit AF488 | Donkey | Invitrogen | A-21206 |

### RT-qPCR

To quantitatively compare the relative abundance of host mRNA, total RNA was extracted using TRIzol reagent (Invitrogen 15596026), purified using Zymo Direct-zol-96 RNA Kit (Zymo R2054). The purified RNA was reverse transcribed using iScript Reverse Transcription Supermix (Bio-rad 1708840). The cDNA was diluted 1:20 using nuclease-free water. The qPCR master mix was prepared using 4 μL of diluted cDNA, 0.5 μL of 10 μM forward and reverse primers (**Table 2**) and 5 μL of PowerTrack SYBR Green master mix (Thermo Fisher. A46109). The qPCR was performed using Applied Biosystems QuantStudio 3 machine. The relative fold change was calculated using cycle threshold (Ct) values following the ΔΔCt method.

**Table 2.**
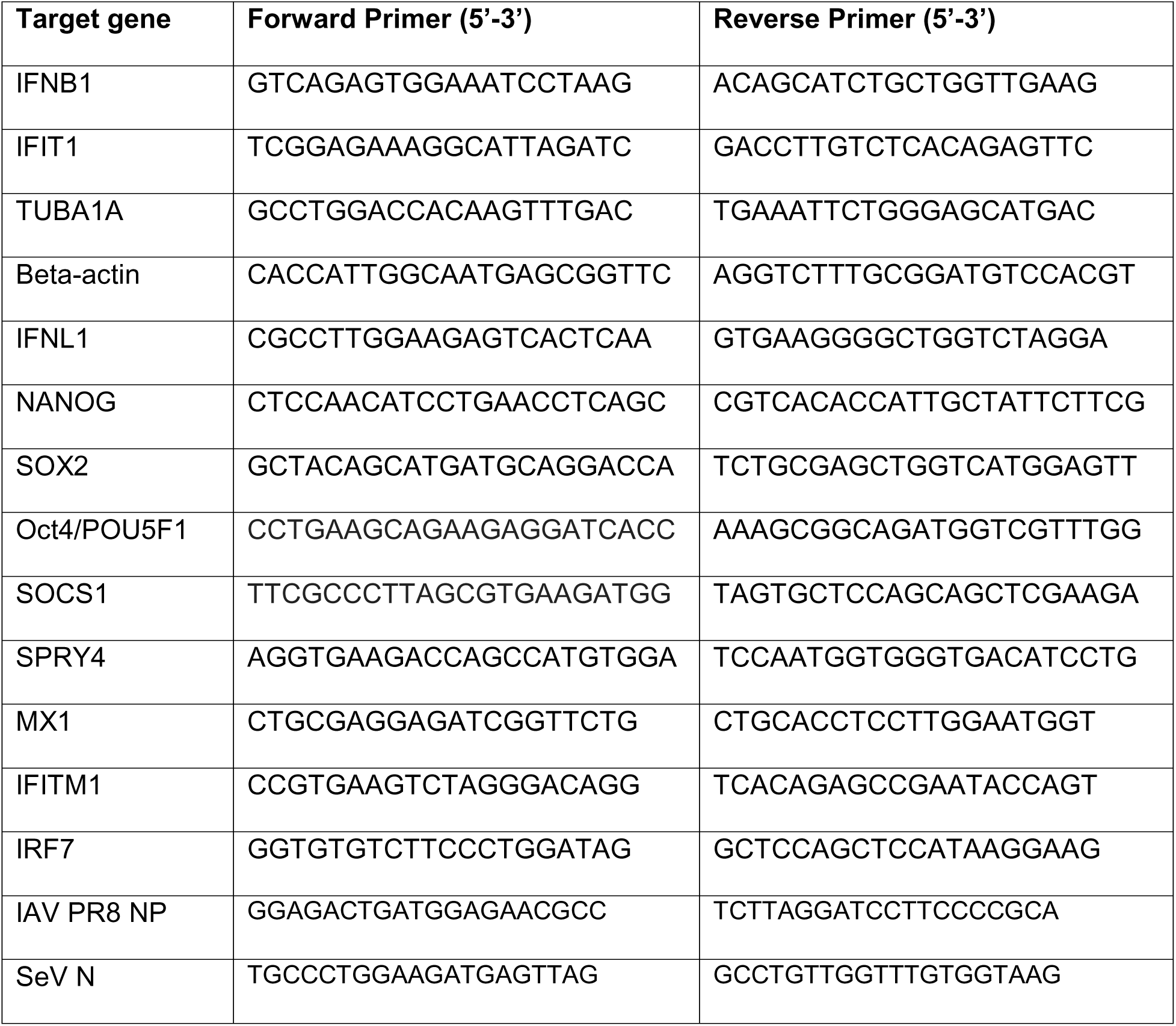
Oligonucleotide sequences for RT-qPCR.

## Data availability

The raw sequencing data sets generated during this study are available on the NCBI Gene Expression Omnibus (GEO) server under accession number GSE319085, GSE318941. All code used for bulk and single-cell RNA-seq analyses are available at https://github.com/BlancoMeloLab/ESC_IAV_RNAseq_analysis.

## ACKNOWLEDGMENTS

This work was funded by the Vaccine and Infectious Disease Division (VIDD) at Fred Hutchinson Cancer Center. D.B-M. is supported by the Kingship Foundation Searle Scholars Program. Q.Y. was supported by the University of Washington Department of Global Health T32 Diseases of Public Health Importance Training Grant (T32AI007509). S.B. is supported by the Deutsche Forschungsgemeinschaft (DFG, German Research Foundation) – 546121141. This research was approved by Fred Hutchinson Cancer Center’s IRB (FHIRB0011124). We thank Dr. Manu Setty (Fred Hutch) for fruitful conversations and useful suggestions on the single cell analyses. We also extend our sincere gratitude to Dr. Benjamin tenOever (NYU) for fruitful conversations, sharing reagents, and for his overall support for this project. We thank Dr. Ram Savan and Dr. Nandan Gokhale for sharing SeV working stocks (ATCC VR-907) and valuable discussions.

