## Supplemental Figures for "Pluripotency Factors Modulate Interferon Signaling in Embryonic Stem Cells"

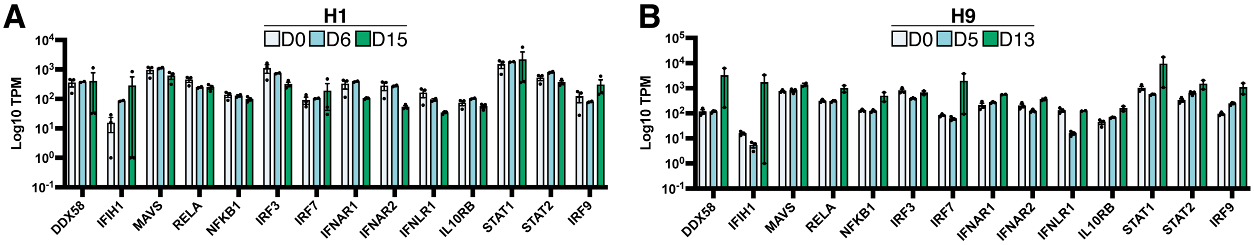


**Fig. S1. Expression levels of IFN effectors in hESC and their corresponding differentiated stages**. (**A** and **B**) expressions of key interferon effectors involved in transcription and response to interferon in H1 (A) and H9 (B) -derived cells are represented by Log10 Transcript per million (TPM). The bar graphs are color-coded based on the cell types. No significant differences were detected through multiple paired t-test (p-value >=0.05).

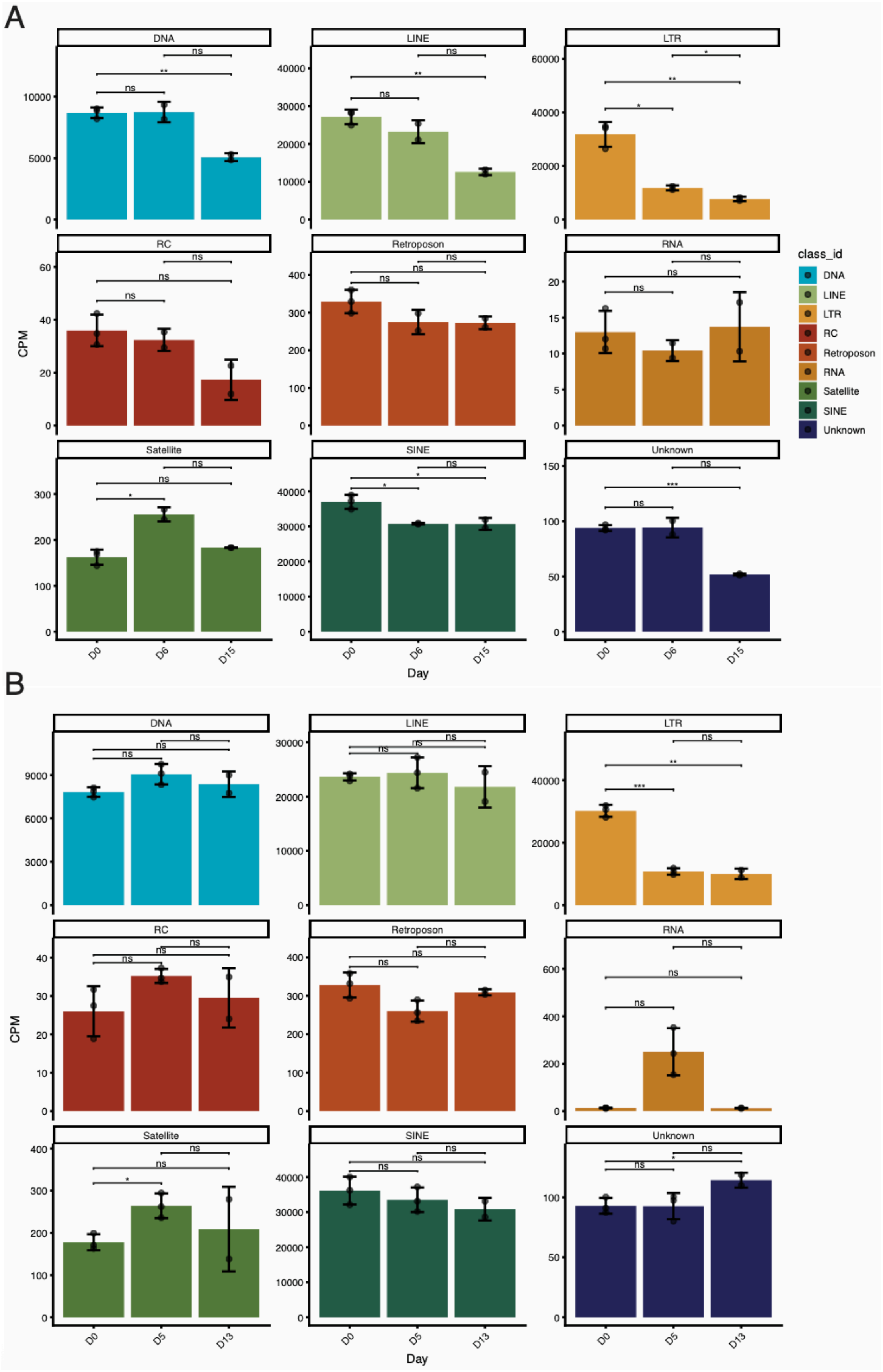
A
DNA
LINE
LTR
RC
Retroposon
RNA
Satellite
SINE
Unknown
CPM
Day
D0
D6
D15
class_id
DNA
LINE
LTR
RC
Retroposon
RNA
Satellite
SINE
Unknown
B
DNA
LINE
LTR
RC
Retroposon
RNA
Satellite
SINE
Unknown
CPM
Day
D0
D5
D13

**Fig. S2. Expression levels of repetitive elements across H1 and H9 differentiation lineage**. (**A** and **B**) Bar plots depicting the normalized counts per million (CPMs) for diverse classes of repetitive elements defined by RepBase (indicated in color, Ref. 52) across differentiation stages in the lung progenitor lineage (A) and the cardiomyocyte lineage (B). LTR: Human Endogenous Retroviruses (HERVs).

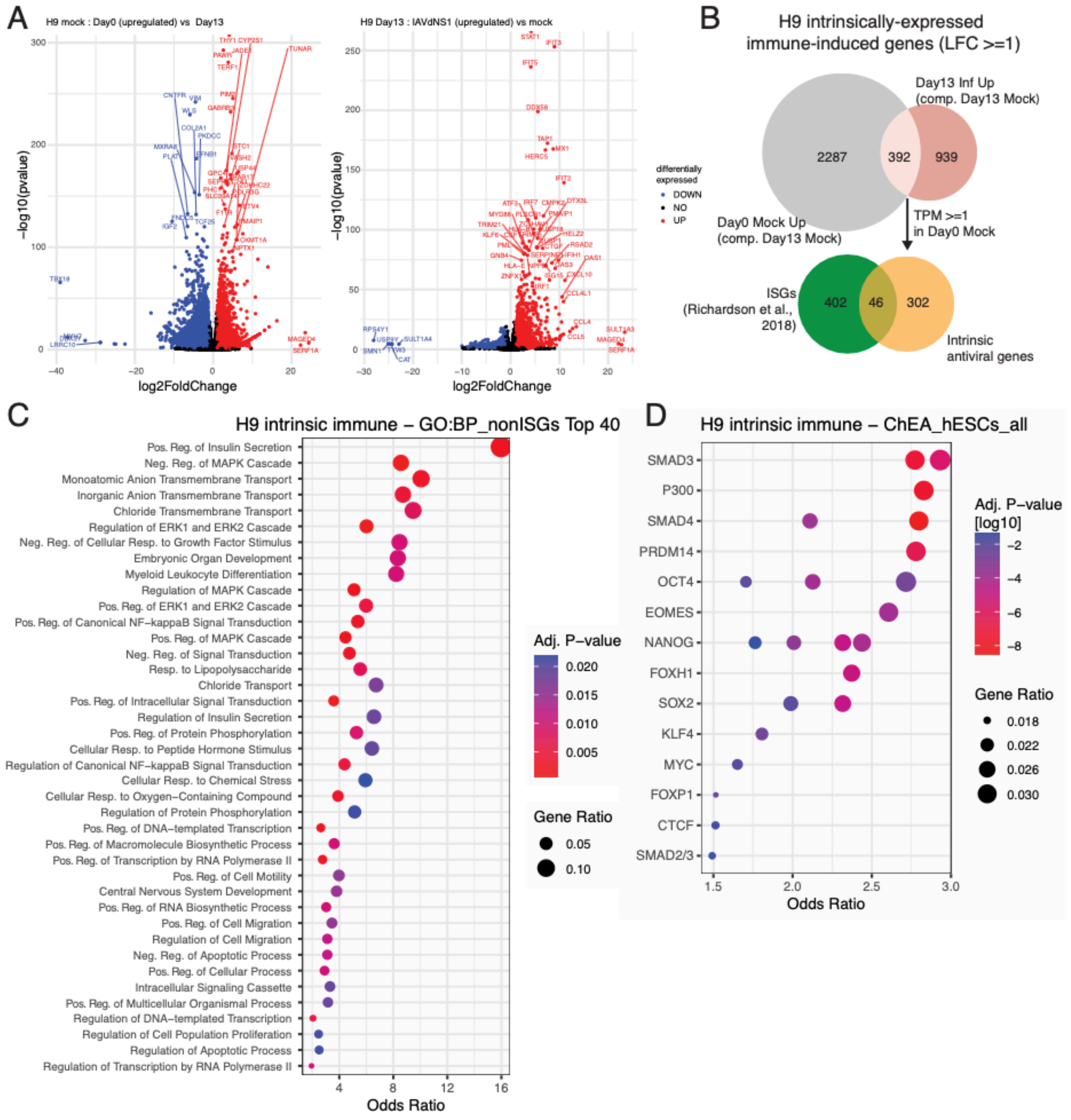


**Fig. S3. Defining intrinsic immune genes in H9 hESCs**. **(A)** Volcano plots highlighting genes upregulated in D0 ESCs compared to D13 differentiated ESCs (left side) and genes upregulated in D13 differentiated ESCs by IAVΔNS1 infection (right side). **(B)** Venn diagram depicting the definition of intrinsically expressed immune genes. **(C)** Top 40 Gene ontology (GO) Biological Processes (BP) terms enriched in intrinsically expressed, non ISG immune genes. **(D)** ChEA 2022 enrichment hits for H9 intrinsically expressed immune genes using ChIP-Seq experiments done in human ESCs.

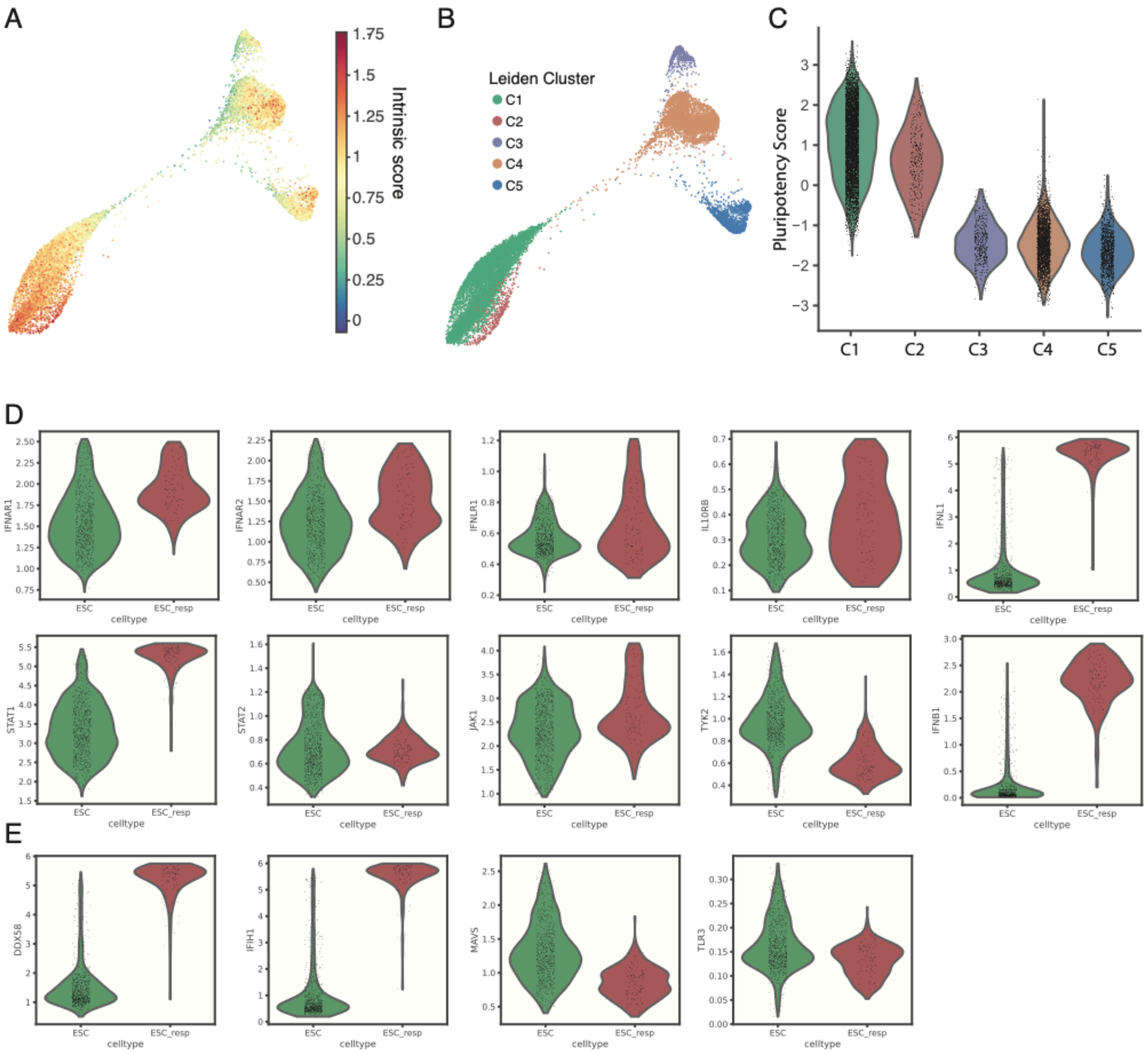


**Figure S4. Force directed layout separates cells from scRNA-seq into distinct clusters that captures transcriptional differences during differentiation and immune activation. (A)** Intrinsic immune gene score across D0 and D6 cells. The list of 712 intrinsic immune genes defined in Fig. 2B was used to calculate an intrinsic immune gene score for each cell. **(B)** Cell annotation according to Leiden clusters based on their transcriptional proximity. **(C)** The average expression scores of pluripotency genes within each Leiden clusters as in (B). The score indicates Z-score across the population, where 0 represents the average expression among all cells and 1 indicates 1 standard deviation from the average. **(D)** Gene expression of proteins involved in IFN pathway. ESC and ESC_resp cells from D0 IAVΔNS1 shown (leiden cluster C1 and C2, respectively). **(E)** Gene expression of pattern recognition receptors (PRRs). ESC and ESC_resp cells from D0 IAVΔNS1 shown (leiden cluster C1 and C2, respectively).

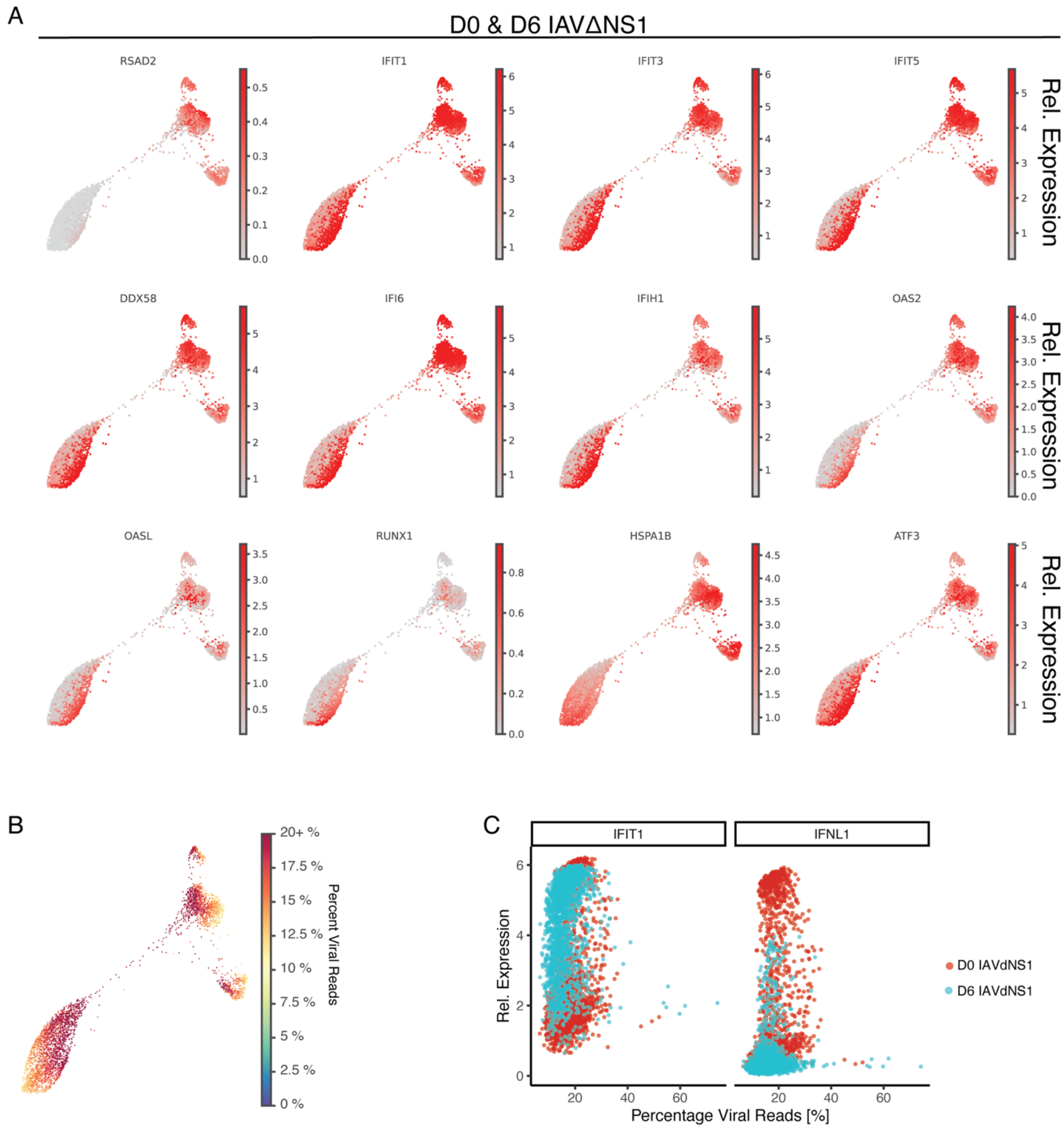


**Figure S5. Expression of ISGs and Viral trasncripts in D0 and D6 IAVΔNS1 infected ESCs.** (**A**) Force-directed layout (FDL) of D0 H1 hESC and differentiated D6 endoderm cells independently infected with IAVΔNS1 at MOI of 3. Relative expression indicates MAGIC-imputed, log-normalized read counts. (**B**) Percent of reads in each cell that correspond to IAV reads. (**C**) Scatter plots depicting the relative expression of IFNL1 and IFIT1 compared to the percent of viral UMI counts for each cell.

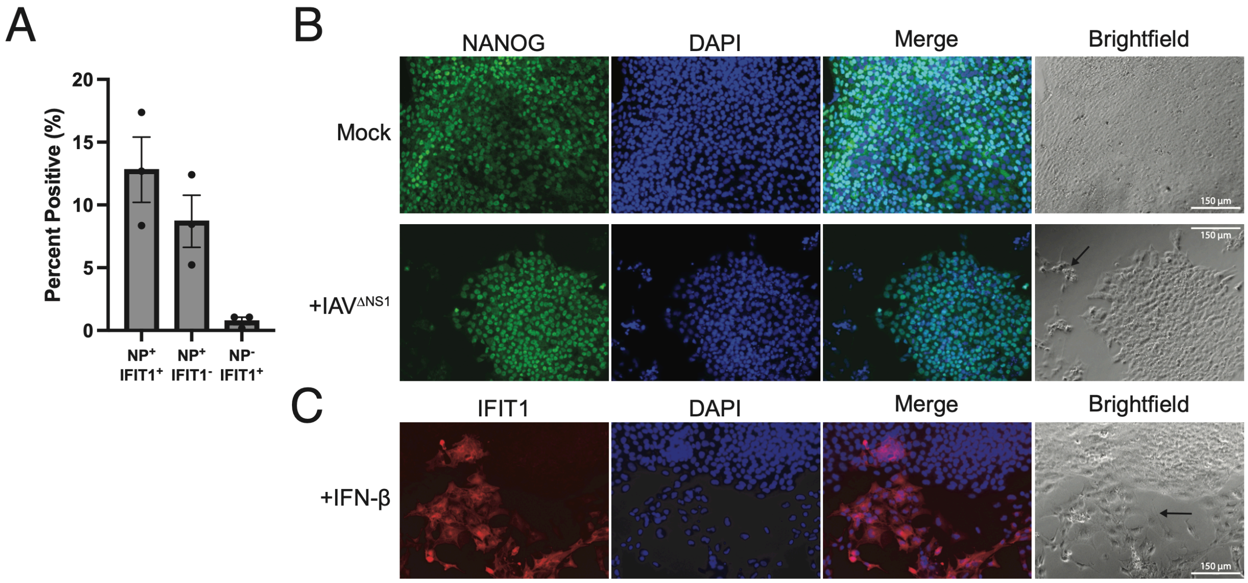


**Figure S6. Immunofluorescence of H1 hESCs validate the stem cell state of IFN-responding cells**. **(A)** Quantification of **Fig. 3G** comparing cells that are expressing viral NP and/or IFIT1. Each data point represents a technical replicate image. **(B)** Comparison of NANOG expression during IAVΔNS1 infection in hESC undergoing spontaneous differentiation, highlighted by arrows. **(C)** Comparison of IFIT expression during IFN-β treatment in hESC undergoing spontaneous differentiation, highlighted by arrows. Scale bar = 150 µm.

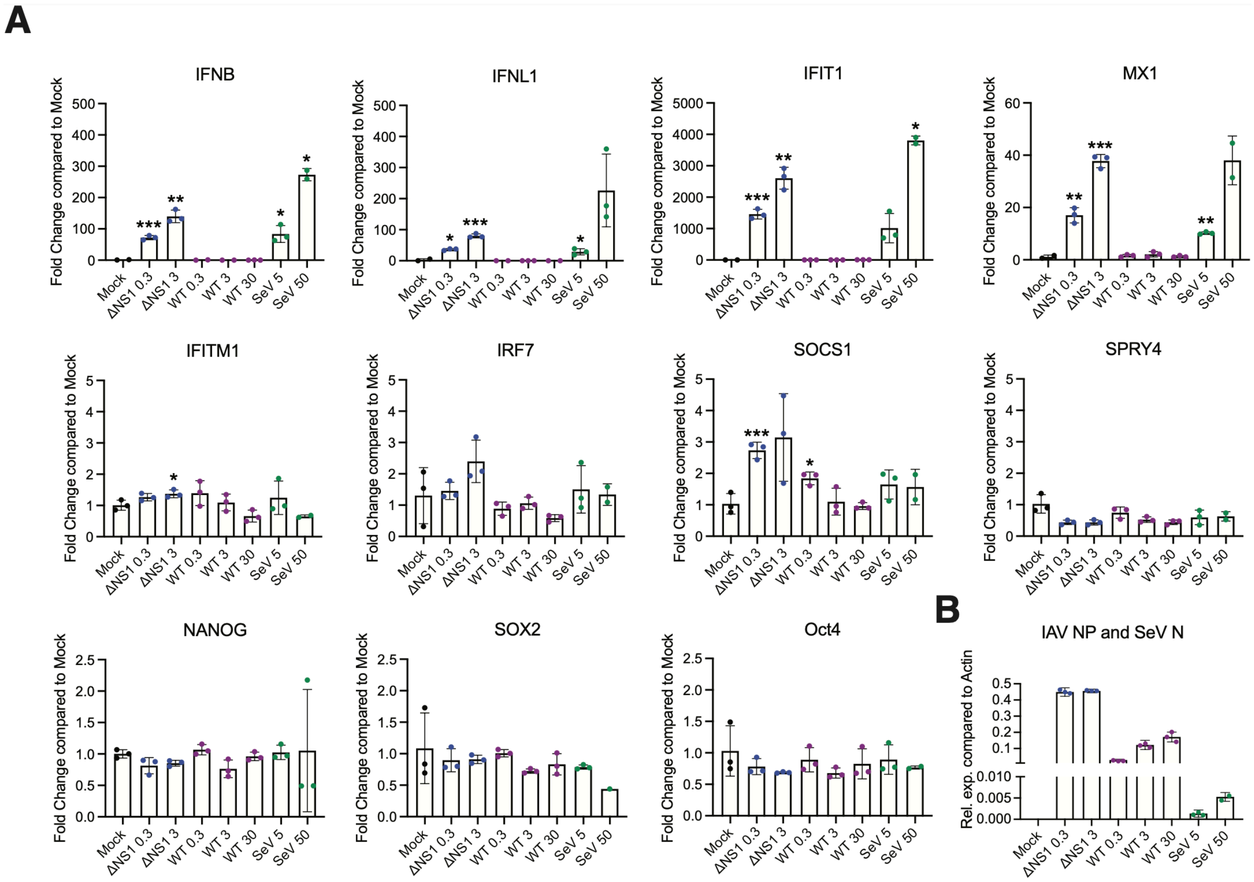


**Figure S7. Multiple virus strains can induce IFN and ISG expression in H1 hESCs.** Gene expression measured by RT-qPCR in H1 hESCs infected with IAVΔNS1, IAV-WT or SeV. Cells were infected for 8 hours at different infectious titers (IAVs: MOI 0.3, 3 and 30; SeV: 5 and 50 HAU/ml) before RNA extraction. Asterisks indicate significance (*p-value < 0.05, ** p-value < 0.01, *** p-value < 0.001) compared to Mock control with Welch's unpaired t-test. **(A)** IFNs, ISGs and pluripotency factors are depicted as fold change compared to mock control. **(B)** Viral genes (NP for IAV and N for SeV) are depicted as dCT compared to beta-actin.

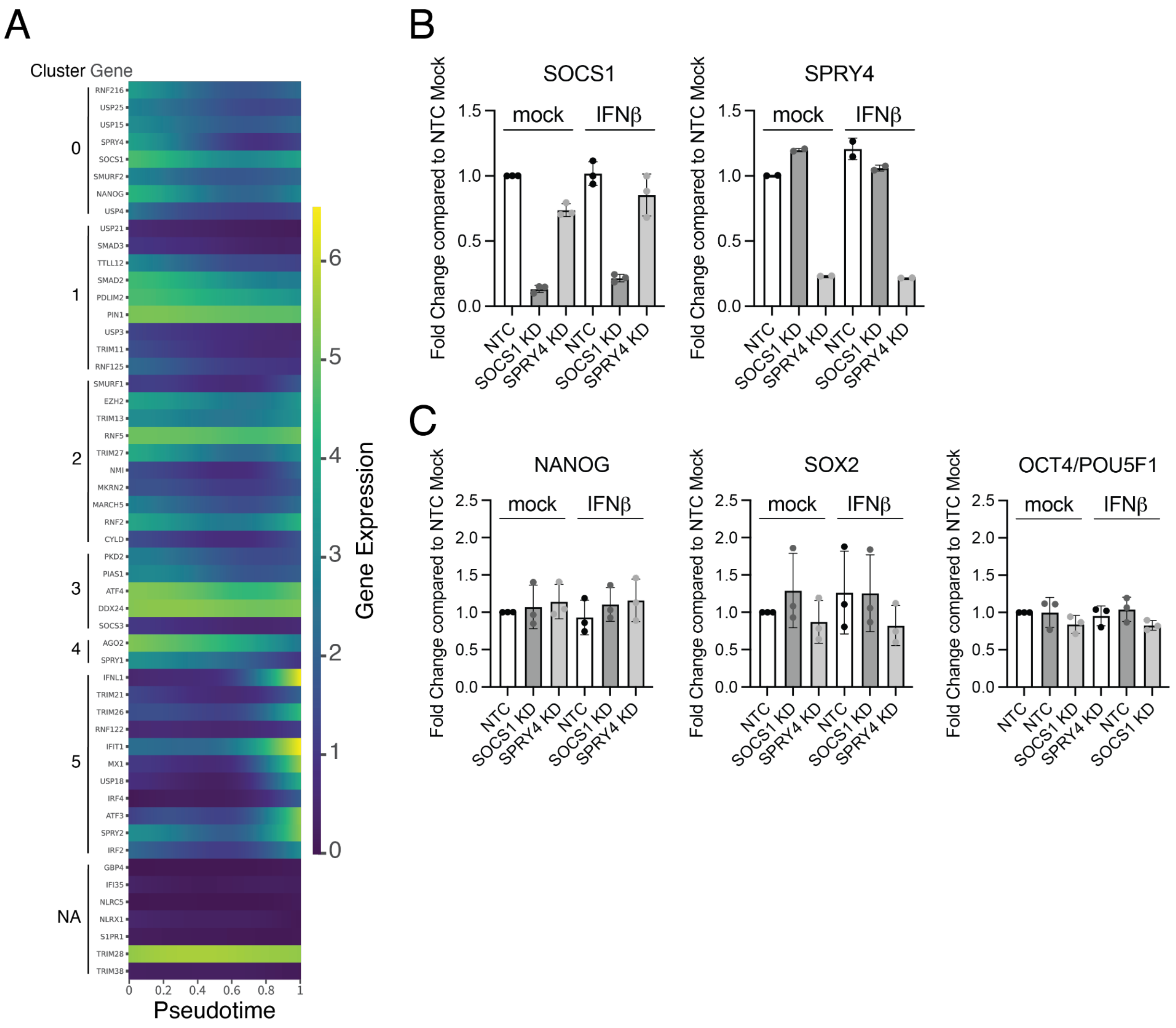


**Figure S8. Negative regulators are expressed in H1 hESCs. (A)** Gene expression of negative regulators of IFN response along pseudotime. Genes shown here are negative regulators as defined by Arimoto et al. (2018), plus NANOG, IFIT1 and MX1. Genes are grouped by clusters defined in Fig. 4C. Genes with cluster NA are not found in any cluster because their variance of gene expression along pseudotime is < 0.01. **(B)** RT-qPCR verification of siRNA-mediated knock-down (KD) in hESCs. **(C)** RT-qPCR quantified expression of pluripotency factors after siRNA-mediated knockdown of SOCS1 or SPRY4. IFNβ treatment at 100 U/ml for 8 hours in H1 hESCs.

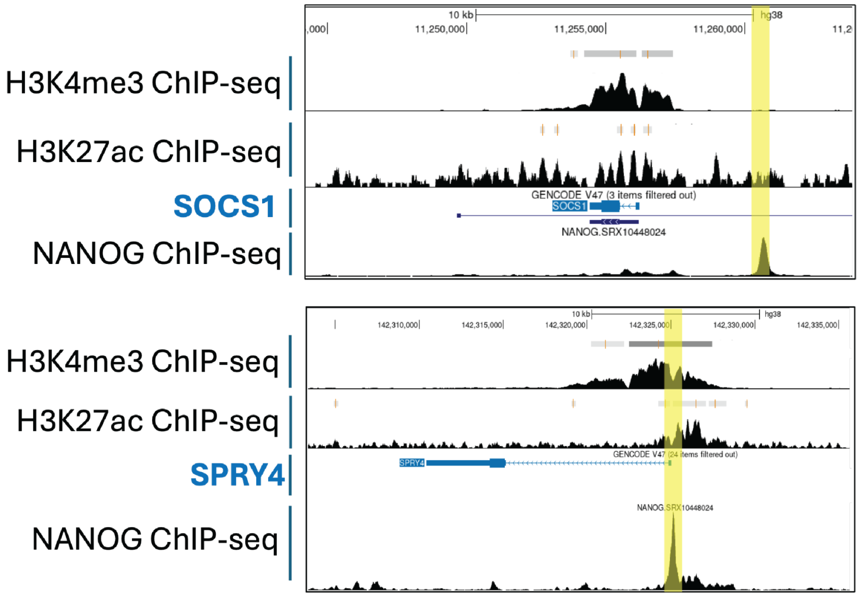


**Figure S9. Negative regulators SOCS1 and SPRY4 are regulated by NANOG in H1 ESCs.** ChIP-Seq peaks for H3K4me3, H3K27ac and NANOG visualized in UCSC Genome Browser for SOCS1 (left side) and SPRY4 (right side) loci in H1 ESCs. ChIP-Seq accession numbers: NANOG (SRX10448024, ChIP Atlas), H3K4me3 (ENCSR814XPE, ENCODE), H3K27ac (ENCSR000ANP, ENCODE).
